# Human Cytomegalovirus induces neuronal enolase to support virally-mediated metabolic remodeling

**DOI:** 10.1101/2022.04.26.488511

**Authors:** Isreal Moreno, Irene Rodríguez-Sánchez, Xenia Schafer, Joshua Munger

**Affiliations:** Department of Biochemistry and Biophysics, School of Medicine and Dentistry, University of Rochester, Rochester, New York, USA; Department of Microbiology & Immunology, School of Medicine and Dentistry, University of Rochester, Rochester, New York, USA

## Abstract

Viruses depend on cellular metabolic resources to supply the energy and biomolecular building blocks necessary for their replication. Human Cytomegalovirus (HCMV), a leading cause of birth defects and morbidity in immunosuppressed individuals, induces numerous metabolic activities that are important for productive infection. However, many of the mechanisms through which these metabolic activities are induced and how they contribute to infection are unclear. We find that HCMV infection of fibroblasts induces a neuronal gene signature, as well as the expression of several metabolic enzyme isoforms that are typically expressed in other tissue types. Of these, the most substantially induced gene was the neuron-specific isoform of enolase (ENO2). Induction of ENO2 expression is important for HCMV-mediated glycolytic activation, as well as for the virally-induced remodeling of pyrimidine-sugar metabolism, which provides the glycosyl subunits necessary for protein glycosylation. Inhibition of ENO2 expression or activity reduced UDP-sugar pools, attenuated the accumulation of viral glycoproteins, and induced the accumulation of non-infectious viral particles. In addition, our data indicate that the induction of ENO2 expression depends on the HCMV U_L_38 protein. Collectively, our data indicate that HCMV infection induces a tissue atypical neuronal glycolytic enzyme to activate glycolysis and UDP-sugar metabolism to provide the glycosyl building blocks necessary for viral protein glycosylation and the production of infectious virions.

**Significance Statement:** Viruses are obligate parasites that obtain energy and mass from their host cell. Control over the metabolic resources of the cell has emerged as an important host-pathogen interaction that can determine infectious outcomes. We find that the Human Cytomegalovirus (HCMV), a major cause of birth defects and morbidity in immunosuppressed patient populations, induces a neuronal gene signature in fibroblasts including the expression of neuronal-specific enolase (ENO2). Our data indicate that ENO2 is important for HCMV-mediated metabolic remodeling including glycolytic activation and the production of pyrimidine sugars, as well as for viral infectivity. These findings indicate that viruses are capable of tapping into alternative tissue-specific metabolic programs to support infection, highlighting an important viral mechanism of metabolic modulation.

## Introduction

Human Cytomegalovirus (HCMV) is a ubiquitous herpesvirus that infects 60-90% of the global population [1]. Characteristic of herpes viruses, HCMV establishes a lifelong latent infection. HCMV infection is asymptomatic in most immunocompetent people; however, severe disease can occur in immunocompromised individuals including cancer patients, AIDS patients, and organ transplant recipients [2]. Additionally, congenital HCMV infection is a leading cause of infectious birth defects including hearing and vision loss, cognitive deficits, and, in severe cases, infant death [3, 4].

Betaherpesviruses are enveloped and characterized by large, double-stranded DNA genomes. HCMV possesses a DNA genome of ∼230 kB that encodes more than 200 open reading frames [5]. The outer viral envelope is studded with viral glycoproteins that are required for host cell recognition, attachment, and fusion [6]. These glycoproteins are diverse, and differentially interact with receptors on a variety of cells types, which contributes to HCMV’s broad cell tropism.

HCMV infection induces substantial changes to the cellular metabolic network [7–9], including inducing glycolysis [8, 10, 11], fatty acid biosynthesis [12, 13], amino acid metabolism [11] and pyrimidine biosynthesis [11, 14]. Inhibiting any of these pathways during viral infection significantly attenuates HCMV production [11–14]. For example, inhibition of *de novo* pyrimidine biosynthesis was found to disrupt the glycosylation of gB leading to protein instability and degradation, and a viral growth defect [14].

HCMV induces glycolysis in several ways. It has been shown that viral infection modulates the activity of key signaling hubs such as the AMP-activated protein kinase (AMPK). AMPK monitors ATP and AMP levels serving as an energy gauge that activates metabolic stress pathways in response to decreased ATP levels [9, 15]. HCMV induces the expression of the high capacity glucose transporter 4 (GLUT4) [16] and activates AMPK [17], both of which have been found to be important for HCMV-mediated increases in glucose consumption [15–17]. More recently, AMPKα2 activity has been found to be necessary to induce GLUT4 expression during HCMV infection [18]. In addition, AMPK directly phosphorylates the carbohydrate-responsive element-binding protein (ChREBP), the levels of which are induced by HCMV infection, and are typically sensitive to glucose concentrations [19]. ChREBP also induces glycolytic enzyme expression such as GLUT4 via promoter binding to the carbohydrate response element (ChoRE) [20].

Here, we find that HCMV infection induces the expression of neuronal genes leading to a global neuronal gene signature. Increased expression of one of these genes, the neuron-specific isoform of enolase, Enolase 2 (ENO2), is necessary for HCMV-induced glycolytic activation, the induction of pyrimidine and pyrimidine sugar metabolism, and high titer viral replication. Relatedly, our results indicate that increased ENO2 expression is important for the accumulation of viral glycoproteins and optimal viral infectivity, highlighting that HCMV induces neuronal enolase to support glycolytic and nucleotide-sugar metabolic remodeling and successful infection.

## Materials and Methods

### Cell Culture and viral infections

Human 293T, telomerase immortalized MRC5 fibroblasts, telomerase immortalized HFF fibroblasts, and all derived cell lines (see below) were cultured in Dulbecco’s modified Eagle medium (DMEM; Invitrogen) supplemented with 10% fetal bovine serum, 4.5 g/liter glucose, and 1% penicillin-streptomycin (Pen-Strep; Life Technologies) at 37°C in a 5% (vol/vol) CO2 atmosphere. Once confluent, cells entered quiescence by culturing in serum-free media for 24h and then infected or mock-infected at the indicated multiplicity of infection (MOI). The viral inoculum was removed after 2h adsorption, cells were rinsed with a sodium citrate inactivation buffer (40 mM sodium citrate, 10 mM KCl and 135 mM NaCl, pH 3.0), and replaced with fresh serum-free media.

The wild-type strain of HCMV used was BAD*wt*, a bacterial artificial chromosome (BAC) clone of AD169 [21]. The TB40/EmCherry-UL99eGFP virus that expresses mCherry under an SV40 promoter and a UL99-GFP fusion protein was a gift from Eain Murphy [22]. HCMV-ΔU_L_38 deletion virus was provided by Thomas Shenk (Princeton University) as a BAC derived virus lacking the entire U_L_38 allele (ADΔU_L_38) [23]. Viral stocks were propagated in MRC5 cells. The titers of the viral stocks were determined either by traditional 50% tissue culture infective dose (TCID_50_) analysis or by calculating Infectious Units per mL (IU/mL). Briefly, IU/mL was calculated by plating 10-fold dilutions of viral stocks in 384-well plates. After incubating for 48 hrs, the cells were fixed with -80°C methanol and stained for IE1 expression. Automated microscopy with a Cytation 5 microscope was used to analyze plates and calculate IU/mL.

### Compounds

Rapamycin (Sigma-Aldrich) and Torin-1 (ApexBio) were prepared at 100uM and 250uM respectively in dimethyl sulfoxide (DMSO). Both were added to cells at 1:1000 dilution to final concentrations of 100 nM and 250 nM respecively. POMHEX was obtained as a gift from Florian Muller [24], but was also ordered from MedChem Express (Cat. No.: HY-131904) and resuspended to 100 mM in DMSO. POMHEX was added to cells to 10-1000 nM final concentration as indicated.

### RNA-seq

Quiescent MRC5 fibroblasts in 10-cm dishes were infected at an MOI of 3.0 with AD169 HCMV. After 48 hpi, TRIzol reagent (Invitrogen) was used to isolate total cellular RNA according to the manufacturer’s instructions. Turbo DNase kit (Invitrogen) was used to DNase treat each sample to remove DNA and subsequently RNA was purified using RNeasy MinElute cleanup kit (Qiagen). A NanoDrop 1000 spectrophotometer was used to quantify total RNA concentrations and RNA quality assessed with an Agilent Bioanalyzer. The next-generation sequencing library construction was made by using the TruSeq stranded mRNA sample preparation kit (Illumina). Briefly, oligo(dT) magnetic beads were utilized to purify mRNA from 200 ng total RNA. Random hexamer priming was used for first-strand cDNA synthesis, followed by second-strand cDNA synthesis using dUTP incorporation for strand marking. Double-stranded cDNA then underwent end repair and 3′ adenylation. cDNA was modified by the ligation of Illumina adapters to both ends. Samples were purified by gel electrophoresis and PCR primers specific to the adapter sequences were used to generate cDNA amplicons ranging in size from 200 to 500 bp. The University of Rochester Genomics Research Center (GRC) prepared the library and performed the sequencing as previously described [25] .

### Infectious virion to DNA ratio

Confluent hTert-HFFs in 15cm plates were infected with HCMV at MOI=1.0, total virus was collected from the cells at 5 dpi. A small aliquot was saved to determine the viral titer by TCID_50_. The remaining was concentrated by sorbitol cushion. Briefly, the viral supernatants were underlaid with a standard sorbitol cushion and centrifuged for 1.5 h at 26,000 rpm. The viral pellet was resuspended in 400 uL of DNase buffer and split into two aliquots. 10 U of DNaseI (10 U/uL) (Invitrogen) was added to one aliquot for 1 hour at 37C. The reaction was stopped by adding 16 uL 25 mM EDTA to a final concentration >2 mM and incubated at 65C for 10 minutes. Total DNA was extracted from both samples by phenol:chloroform:IAA (25:24:1) extraction. Viral DNA abundance was determined by RT-qPCR using IE1 specific primers. A preparation of BAC*wt* DNA was used to generate a standard curve to quantify genome copy numbers.

### shRNA Knockdown and rescue

To achieve targeted knockdown of ENO2 in HFFs, a pLKO.1-based Mission shRNA construct targeting ENO2 (Sigma/Broad Institute clone number TRCN0000157687) obtained from the Sigma-Aldrich Mission shRNA library was selected after a screening process in which ENO2-knockdown was assessed by qPCR and western blot. Mission pLKO.1-puro control construct (Sigma SHC001) was used as a control in all shRNA experiments.

To generate an shRNA-resistant cDNA clone of ENO2, ENO2 was amplified from cDNA obtained from fibroblasts cells with homologous overhangs for cloning into pLenti CMV/TO Blast via Gibson Assembly (Forward 5’-GAACCAATTCAGTCGACTGGATCCATGTCCATAGAGAAGATCTGGG-3’, Reverse 5’-ACCACTTTGTACAAGAAAGCTGGGTCTAGATCACAGCACACTGGGATTACGGAAGTT-3’). Site-directed mutagenesis was then used to introduce four silent point mutations at the site targeted by the shRNA construct (CTCAAGGGAGTCATCAAG to CTCAA**A**GG**C**GT**T**AT**T**AAG) using the following primers Forward 5’-ACACTCAAAGGCGTTATTAAGGACAAATACGGCA-3’, Reverse 5’-GTCCTTAATAACGCCTTTGAGTGTATGGTAGACC-3’. This shRNA-resistant cDNA variant of ENO2 was then cloned into XbaI/BamHI digested pLenti CMV/TO Blast plasmid (Addgene plasmid #17486) by Gibson Assembly following manufacturer’s instructions. All plasmid preps and lentiviral transductions were performed according to the manufacturer’s instructions.

### Lentiviral transduction and cell line generation

Pseudotyped lentivirus was produced in 293T cells seeded in 10-cm dishes at a density of 2×10^6^ cells per cm^2^ and grown for 24 hr prior to transfection with 2.6 μg lentiviral vector, 2.4 ug PAX2, and 0.25 ug vesicular stomatitis virus G glycoprotein expression plasmid using FuGENE 6 (Promega) or TransDuceIT (Mirus). The medium was aspirated after 24 hr, and 4 ml fresh medium was added to the plate. After another 24 hr, the supernatant was filtered through a 0.45-um filter and stored at -80°C. The thawed supernatant was applied to fibroblast cells in the presence of 5 ug/ml Polybrene (Millipore). The inoculum was incubated for 24 hr, after which the medium was then refreshed and the cells allowed to recover for 72 hr prior to selection with 1 ug/mL puromycin (VWR). To rescue shRNA knockdown, a second round of lentiviral transduction procedure was repeated by treating the shRNA-treated, puromycin-selected cells with the shRNA-resistant ENO2 construct. After treatment, these cells were selected with 5 ug/mL blasticidin.

ENO2 knockout and control cell lines were generated in HFFs and MRC5 fibroblasts using CRISPR-Cas9 ribonucleoproteins (RNP) system. Guide RNA targeting ENO2 (ggugaaggaagccaucgaca) was designed and ordered from Synthego. Guide RNA and tracrRNA (Synthego) were then duplexed. Duplexed RNA and Cas9 were incubated together to generate RNPs. HFF-hTs were prepared following manufacturer’s instructions and RNPs were then electroporated into cells using the Neon electroporator set to 1650 volts, 10 ms pulse width, 3 pulses. Cells were then plated in one well of a 12-well plate. Cas9 lacking duplexed RNA was electroporated as a negative control. Cells were allowed to fill in and expanded. To determine knockdown efficiency, DNA was harvested and ENO2 specific primers were used to amplify the target region. The amplicon was purified and sequenced by Sanger Sequencing (Genewiz). Knockout efficiency was determined to be 82% by Synthego’s Inference of CRISPR Edits (ICE) analysis software (https://ice.synthego.com/#/) to deconvolute sequencing reads (Fig S3).

### Quantification of Metabolite Concentrations

For quantification of lactate secretion and glucose consumption, cells were plated in 10-cm dishes. Once confluent, the medium was removed and serum-free medium was added. An aliquot of this serum-free virgin medium was saved to be used as t = 0 control. Cells were maintained in serum-free medium for 24h, at which time the conditioned medium was collected for glucose measurement or LC-MS/MS analysis to quantify lactate [11]. In addition, cells were harvested in RIPA buffer (Tris-HCl, 50 mM, pH 7.4; 1% Triton X-100; 0.25% Na-deoxycholate; 150 mM NaCl; 1 mM EDTA) buffer for western blot or trypsinized for total cell counts. For quantification of lactate levels, conditioned medium samples were first diluted 1/2 in OmniSolv Water and subsequently diluted 1/100 in 80% methanol. One hundred ul of the methanol diluted samples were centrifuged at 4°C for 5 minutes at full speed to pellet insoluble material. Lactate levels were measured by LC-MS/MS as indicated below.

Glucose levels were quantified using the HemoCue Glucose 201 System (HemoCue) [11]. A glucose standard curve was utilized for each experiment using the t = 0 virgin DMEM medium (4.5 g/liter glucose) serially diluted in PBS. Conditioned medium samples were then diluted serially 1/4 in PBS to ensure signal linearity. The amount of glucose present in each sample was measured using the HemoCue System and normalized using the generated standard curve. To obtain consumption values, the glucose value measured for normalized virgin DMEM medium was subtracted from the result of each normalized conditioned medium value. These values were then normalized to the number of live cells counted in each plate.

For relative quantification of intracellular metabolite concentrations, cells were plated in 10-cm dishes, and once confluent, the medium was removed and changed to serum-free medium. Cells were maintained in this serum-free medium for 24h and then infected with AD169 as described above. 24 hrs prior to harvesting, medium was changed to serum-free medium supplemented with 10 mM HEPES, 1% penicillin-streptomycin. One hour prior to metabolite extraction medium was once again changed. The medium was aspirated and 80:20 OmniSolv Methanol: OmniSolv Water (80% methanol) at −80°C was immediately added to quench metabolic activity and extract metabolites. Cells were then incubated at −80°C for 10 min. Following cell quenching, cells were scraped in the dish and kept on dry ice, and the resulting cell suspension vortexed, centrifuged at 3,000 × g for 5 min, and re-extracted twice more with 80% methanol at −80°C. The three extractions were pooled and completely dried under N2 gas, dissolved in 200 μl 50:50 OmniSolv Methanol: OmniSolv Water methanol, and centrifuged at 13,000 × g for 5 min to remove debris. Samples were analyzed by LC-MS/MS for as indicated below. Relative intracellular metabolite levels were determined by normalizing the peak heights by the total live cells counted for each sample.

For carbon tracing, cells were prepared and HCMV infected as described above. At 48 hpi, media was swapped for either 4.5 g/L U-^13^C-Glucose (Cambridge Isotope Laboratories) or unlabeled glucose. Samples were harvested and processed as indicated above after the indicated time of labeling (0, 30, 60, or 120 minutes). To quantify UDP-Glucose or UDP-GlcNac levels, the concentrations of each metabolite in unlabeled samples was estimated by the comparison of each SRM peak heights to those of a standard dilution curve. Estimated values were then normalized to the number of live cells in each sample. The rate of ^13^C labeling for UDP-Glucose and UDP-GlcNac were determined by multiplying the ratio for the specific labeled species at each time point by the total concentration of either metabolite at T=0.

### LC-MS/MS analysis

As previous described, samples were analyzed using reverse-phase chromatography with an ion-paring reagent in a Shimadzu HPLC coupled to a Thermo Quantum triple quadrupole mass spectrometer running in negative mode with selected-reaction monitoring (SRM) specific scans. The publicly available mzRock machine learning toolkit (http://code.google.com/p/mzrock/) was used to analyze the LC-MS/MS data. The mzRock allows for the automation of SRM/HPLC detection, grouping, signal-to-noise classification, and comparison to known metabolite retention times [26].

### Immunoblotting

Cellular extracts were harvested by scraping in disruption buffer (50 mM Tris [pH 7.0], 2% SDS, 5% 2-mercaptoethanol, and 2.75% sucrose) for Western analysis. Samples were sonicated, boiled for 5 minutes, and centrifuged at 14,000 x g for 5 minutes to pellet insoluble debris. The samples were then separated on a 4-20% SDS-PAGE gradient gel and transferred to a PVDF membrane using the iBlot transfer system. The membranes were blocked by incubation in 5% milk in TBST (50 mM Tris-HCl, pH 7.6, 150 mM NaCl, 0.1% Tween 20), and reacted with primary and, subsequently, secondary antibodies. Protein bands were visualized using an enhanced chemiluminescence (ECL) system (Bio-Rad) and by using the Molecular Imager Gel Doc XR+ system (Bio-Rad). The primary antibodies used were specific for glyceraldehyde-3-phosphate (GAPDH; Cell Signaling), Enolase 2 (ENO2; Santa Cruz Biotechnology), Enolase 1 (ENO1; Santa Cruz Biotechnology), U_L_26 (7H19), U_L_38 (8D6), IE1, pp28 (10B4-29 [27]), HCMV glycoprotein B (gB; Santa Cruz Biotechnology), tubulin (Tub; Cell Signaling), p70 S6 Kinase (S6K; Cell Signaling), phospho-p70 S6 Kinase (Thr389) (pS6K; Cell Signaling).

### Real-time qPCR

Total cellular RNA was extracted with TRIzol (Invitrogen) and used to generate cDNA using SuperScript II reverse transcriptase (Invitrogen) according to the manufacturer’s instructions. Transcript abundance was measured by real-time PCR analysis using Fast SYBR green master mix (Applied Biosystems), a model 7500 Fast real-time PCR system (Applied Biosystems), and the Fast 7500 software (Applied Biosystems). Gene expression equivalent values were determined using the 2^-ΔΔ^*^CT^* method and normalized to GAPDH levels. The following primers were used for real-time PCR: ENO1 5’-GCCTCCTGCTCAAAGTCAAC-3’ (Forward) and 5’-AACGATGAGACACCATGACG-3’ (Reverse); ENO2 5’-AGCCTCTACGGGCATCTATGA-3’ (Forward) and 5’-TTCTCAGTCCCATCCAACTCC-3’ (Reverse); GAPDH 5’-CAT GTT CGT CAT GGG TGT GAA CCA-3’ (Forward) and 5’- ATG GCA TGG ACT GTG GTC ATG AGT-3’ (Reverse); IE1 5’-TGATTCTATGCCGCACCATGTCCA-3’ (Forward) and 5’-AGAGTTGGCCGAAGAATCCCTCAA-3’ (Reverse); HCMV glycoprotein B 5’- TAGCTACGACGAAACGTCAAAA-3’ (Forward) and 5’-GGTACGGATCTTATTCGCTTTG-3’ (Reverse).

### Statistical Analysis

Statistical analyses of the data were performed using Graphpad Prism 9™ statistical software. Statistics used are as indicated and the Tukey test was used to determine significance after ANOVA. For mass spectrometry data analysis, cell count normalized peak heights were analyzed using the publicly available software MetaboAnalyst 3.0 (http://www.metaboanalyst.ca) [28]. The data were auto-scaled, i.e., mean-centered and divided by standard deviation of each variable, followed by PCA multivariate analysis and hierarchical clustering. Statistical significance was determined by one-way ANOVA. Post-hoc analysis was performed using the Fisher’s least significant difference method (Fisher’s LSD).

## Results

### HCMV infection strongly induces the expression of neuronal enolase, which is part of a broader neuronal gene expression signature

HCMV infection markedly impacts host gene expression, including inducing the expression of numerous metabolic enzymes [8, 10]. To further explore the impact of HCMV infection on cellular gene expression, we performed RNA-seq on mock and HCMV-infected fibroblasts at 48 hpi. Of the genes whose expression levels were significantly induced, several were neuronal (Fig 1A), including the neuronal isoform of enolase (ENO2). Ontology analysis of the significantly induced genes indicated that six of the top 10 GO Biological Processes were neuronally related (Fig 1B). As previously described [8], HCMV infection substantially induced the expression of several metabolic enzymes (Fig 1C). In addition to ENO2, there were several tissue atypical isoforms induced by HCMV infection (Fig 1C), including the platelet-specific phosphofructokinase (PFKP), and the skeletal muscle hexokinase (HK2) (Fig 1C). To further investigate how infection impacts the accumulation of ENO2, we analyzed its expression over a time course of infection. HCMV infection substantially increased ENO2 RNA accumulation at every time point analyzed (Fig 1D). Similarly, ENO2 protein accumulated over the same time period during viral infection, but was not detected during mock infection (Fig 1E). These data indicate that HCMV strongly induces ENO2 expression in fibroblasts.

**Figure 1.**
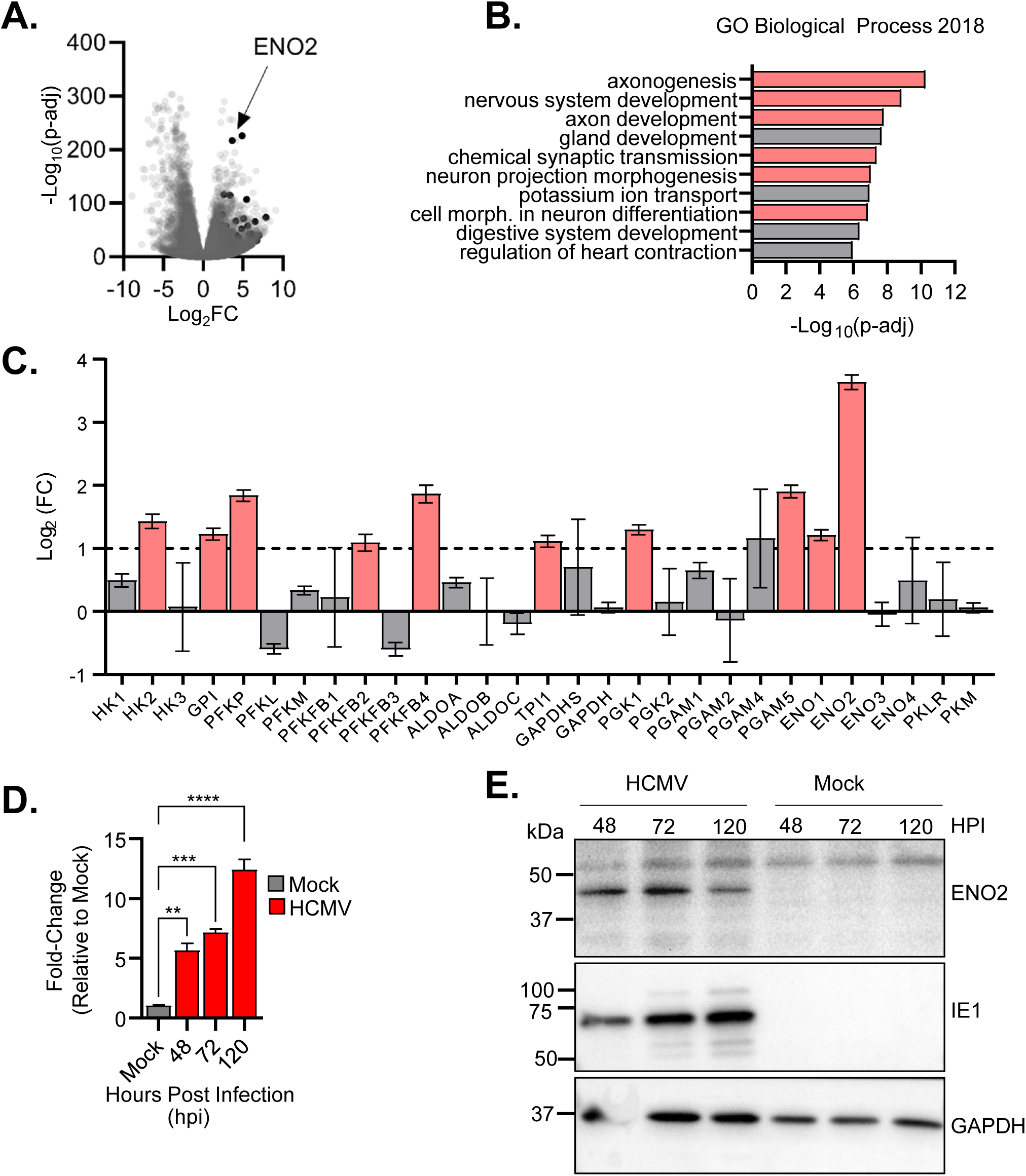
Neuronal Enolase 2 is upregulated by HCMV infection. (A-C) MRC-5 fibroblasts cells were mock or infected with AD169 (MOI 3.0) and total RNA harvested at 48 hpi. RNA-seq analysis was performed. (A) Volcano plot of all genes with a p-value < 0.05, in black are all neuronal related genes identified in (B). (B) GO term analysis was performed using genes that were >4-fold upregulated (Log_2_(FC)≥2). Highlighted in red are all GO terms related to neurological processes. (C) All isoforms of glycolytic genes are shown with Log_2_(FC)≥1 and p-values<0.05 highlighted in red. Values are [mean] ± SEM. (D,E) HFF-hT fibroblasts were mock or AD169 infected (MOI 3.0) and harvested at indicated time points. (D) RT-qPCR performed using ENO2 or GAPDH specific primers. Signals normalized to GAPDH levels and compared to mock infected cells. (E) Protein levels of ENO2 were determined by SDS-PAGE and Western analysis. Values are mean ± SEM (N = 3) **, P< 0.01; ***, P<0.001; ****, P<0.0001.

### Induction of Enolase 2 expression is critical for HCMV-mediated activation of glycolysis

To investigate the role of Enolase 2 during infection, we targeted ENO2 transcripts with shRNA (Fig 2). ENO2 knockdown was confirmed by qPCR (Fig 2A) and western blotting analysis (Fig 2B). To investigate the metabolic impact of targeting ENO2 with shRNA, we profiled metabolite pools via LC/MS-MS (Tables S1 & S2, Fig S1). PCA analysis segregated all of the different sample populations from each other, with the differences between metabolite levels in mock and HCMV infection responsible for the bulk of the data variation in PC 1 (Fig 2C). Transduction with shENO2 induced a substantial metabolic shift in HCMV infected cells as illustrated by PC 2 (Fig 2C). Mock-infected cells transduced with shENO2 largely overlapped with the EV controls as would be expected given the lack of ENO2 expression in fibroblasts (Fig 2C). During infection, ENO2 knockdown increased the levels of fructose-6-phosphate and fructose bisphosphate (Fig 2D & 2E). In contrast, ENO2 knockdown decreased the levels of phosphoenolpyruvate and 3-phosphoglycerate, glycolytic metabolites further downstream in the pathway (Fig 2D & 2E). These results suggest that inhibition of HCMV-mediated induction of ENO2 expression could be blocking virally-activated glycolysis, resulting in increased upstream glycolytic pools and decreased downstream glycolytic pools (Fig 2D). To test this possibility, we analyzed the impact of shENO2 treatment on glucose consumption and lactate excretion. As has been previously reported [11, 12], HCMV infection substantially induced glucose consumption and lactate excretion (Fig 2F & 2G). Knockdown of ENO2 largely prevented the induction of both glucose consumption and lactate excretion (Fig 2F & 2G). Notably, the inhibitory impact of shENO2 on glycolysis, was specific for HCMV-infected cells (Fig 2F & 2G). In mock-infected cells, the levels of glycolytic metabolites and activity were not impacted by shENO2 transduction, which would be predicted given that ENO2 is not substantively expressed in uninfected fibroblasts (Fig 2B). Collectively, our data indicate that HCMV-mediated glycolytic activation depends on the viral induction of ENO2.

**Figure 2.**
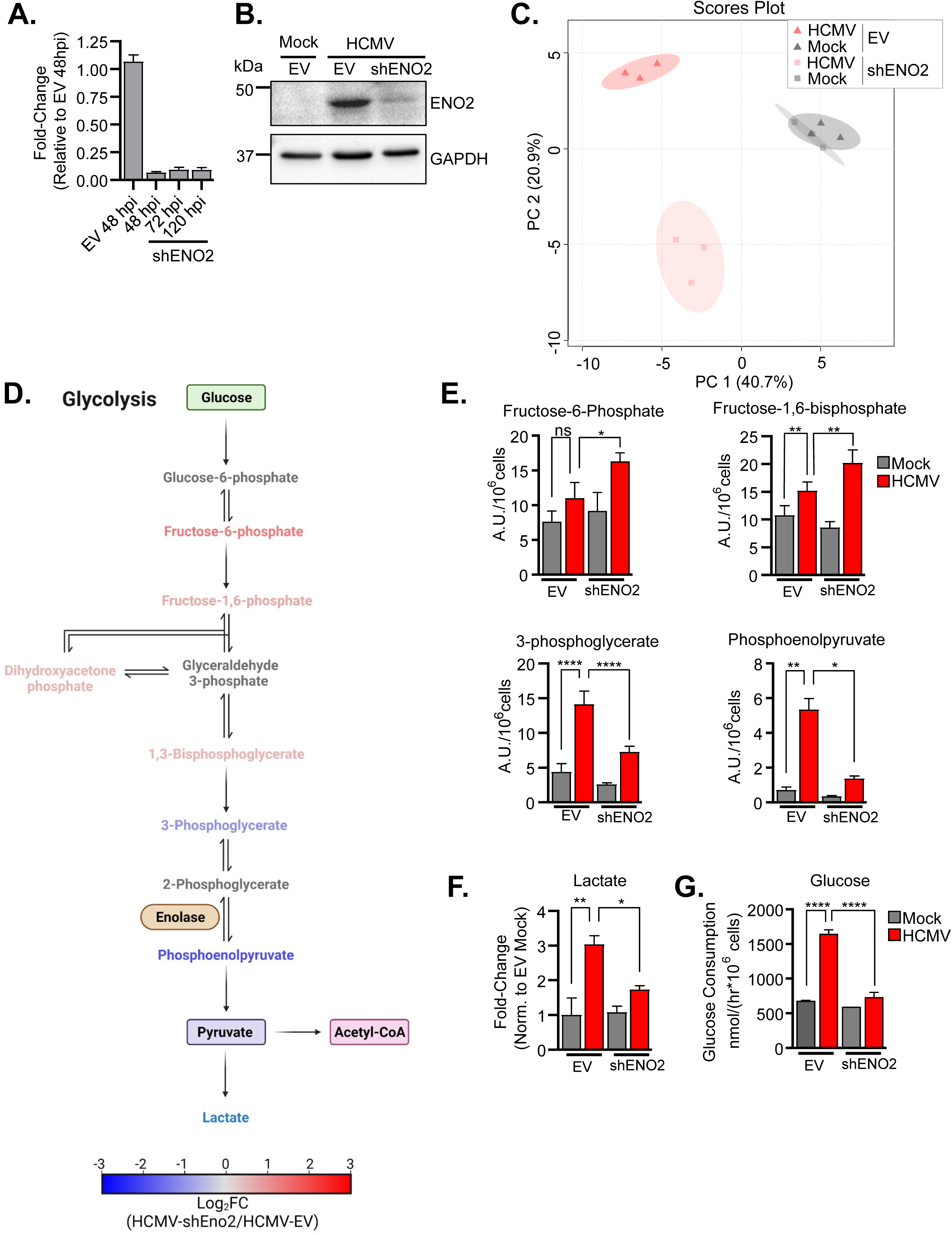
Induction of ENO2 expression is important for HCMV-induced metabolic changes. (A-B) HFF-hT fibroblasts were treated with shRNA targeting Enolase 2 (shENO2) or pLKO-empty vector control (EV) and either Mock or infected with AD169 (MOI 3.0). (A) RNA abundance of ENO2 was determined at indicated time points post infection by RT-qPCR using ENO2 or GAPDH specific primers. Signals normalized to GAPDH levels and compared to EV control cells 48 hpi. (B) Representative western blot of ENO2 protein levels 48 hpi. (C-F) Cells were treated and infected as in A & B. At 48 hpi, metabolic activity was quenched, metabolites extracted, and MS analysis performed. (D) Schematic of glycolysis summarizing the metabolic changes observed during viral infection of shENO2 treated cells as compared to EV. (E) Metabolites in glycolysis are shown. (G-F) Cells were treated as in A & B. Cellular medium was refreshed at 36 hpi and harvested at 60 hpi (24 hours). (F) Lactate abundance was determined by MS analysis. (G) Glucose concentrations were determined. Values are mean ± SEM (N = 3). ns, not significant; *, P<0.05; **, P< 0.01; ****, P<0.0001.

### Induction of ENO2 is important for HCMV-mediated increases in pyrimidine and pyrimidine-sugar metabolites

Glucose-derived carbon feeds the pentose phosphate pathway, which provides the ribose necessary for nucleotide biosynthesis (Fig 3A & 3B). Many of the metabolites in the pentose phosphate pathway were induced by HCMV infection, and notably, substantially further induced upon shENO2 transduction (Fig 3A & 3B), including large increases in 6-phospho-gluconate, ribose phosphate, and sedeoheptulose-7-phosphate. While these pentose phosphate metabolites were increased by shENO2 treatment, the levels of pyrimidine biosynthetic metabolites, pyrimidines, and pyrimidine sugars were generally decreased by transduction with shENO2 (Fig 3A & 3B). These decreased pools include N-carbamoyl-L-aspartate, whose production is the rate-determining step of pyrimidine biosynthesis, as well as UTP, and UDP-GlcNac, which serves as the glycosyl donor for O-linked glycosylation reactions. Notably, in mock-infected cells, the UDP-GlcNac pool was also decreased upon shENO2 transduction cells, although the levels of N-carbamoyl-L-aspartate and UTP were not dramatically impacted. Taken together, these data suggest that induction of ENO2 expression is important for maintaining pyrimidine and pyrimidine sugar pools during HCMV infection.

**Figure 3.**
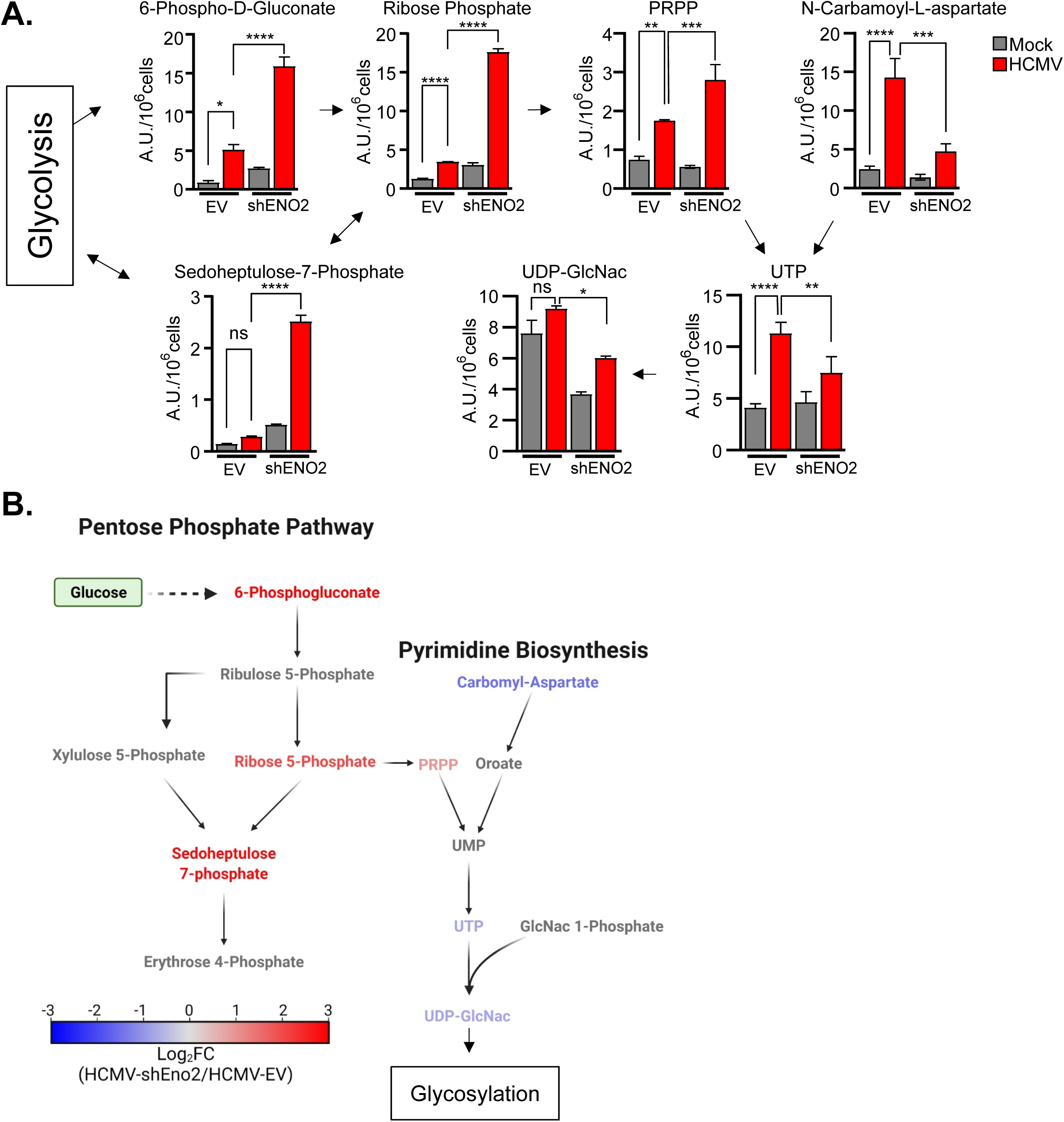
The impact of shENO2 targeting on the pentose phosphate and pyrimidine biosynthetic pathways. HFF-hT fibroblasts were treated with shRNA targeting Enolase 2 (shENO2) or pLKO-empty vector control (EV) and either Mock or infected with AD169 (MOI 3.0). At 48 hpi, metabolic activity was quenched, metabolites extracted, and MS analysis performed. Key metabolites of the pentose phosphate branch point are shown in (A) and summary schematic shown in (B). Values are mean ± SEM (N = 3). ns, not significant; *, P<0.05; **, P< 0.01; ***, P<0.001; ****, P<0.0001.

### The HCMV U_L_38 gene is important for the induction of ENO2

Previously, we have found that the U_L_38 gene is important for HCMV-mediated activation of glycolysis [11], raising the possibility that U_L_38 might contribute to the induction of ENO2 expression. To explore this possibility we analyzed the accumulation of ENO2 in cells infected with WT or a U_L_38 deletion mutant (Fig 4A). ENO2 proteins accumulate substantially less in cells infected with the ADΔU_L_38 mutant relative to those infected with WT AD169 (Fig 4A). Given that U_L_38 induces mTOR activity [11, 29], we sought to determine if mTOR activation was important for HCMV-mediated induction of ENO2. Treatment with rapamycin, an mTORC1 inhibitor, or Torin-1, an ATP analog that inhibits both mTORC1 and mTORC2, significantly inhibited the phosphorylation of S6K, an mTORC1 target [30] (Fig 4B). Treatment with rapamycin had little impact on ENO2 accumulation during infection, whereas Torin treatment significantly reduced the accumulation of ENO2 (Fig 4B). These data suggest that the HCMV U_L_38 protein and mTORC2 play an important role in the HCMV-mediated induction of ENO2. Similar results were observed when tested in MRC-5 fibroblasts (Fig S2).

**Figure 4.**
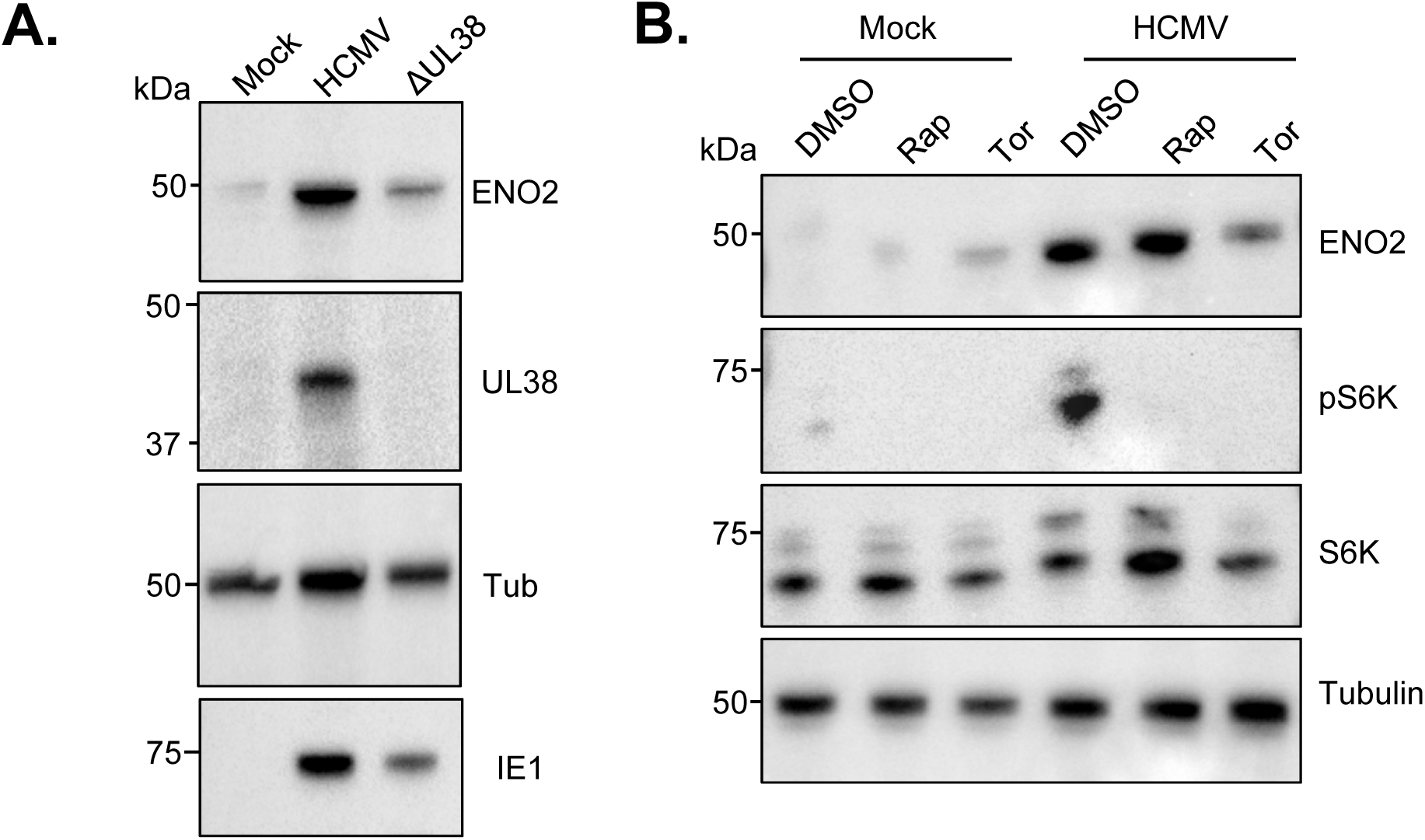
The HCMV protein U_L_38 and mTOR activation are required for HCMV-mediated induction of ENO2 expression. (A) HFF-hT fibroblasts were either mock, AD169 infected (MOI 3.0), or AD169ΔU_L_38 infected (MOI 3.0). Total protein was collected at 48 hpi. SDS-PAGE and Western blot analysis performed with the indicated antibodies. (B) HFF-hT were either mock or AD169 infected (MOI 3.0) and treated with mTOR inhibitors Rapamycin (Rap) or Torin (Tor) or DMSO as a control. Total protein was collected at 48 hpi. SDS-PAGE and Western blot analysis performed with the indicated antibodies.

### Induction of Enolase 2 is important for HCMV infection

To explore how the induction of ENO2 contributes to productive HCMV infection, we utilized an shRNA-resistant ENO2 expression construct to rescue the expression of ENO2 upon shRNA knockdown. While this rescue construct did not fully restore ENO2 expression at the latest time points of infection, it substantially restored ENO2 induction upon shRNA-mediated knockdown (Fig 5A). Subsequent analysis of the production of infectious viral progeny indicated that shENO2 knockdown inhibited viral replication by ∼8-fold, which was largely restored by the expression of shRNA-resistant ENO2 (Fig 5B). Further, re-expression of ENO2 substantially rescued the reduced glucose consumption observed after ENO2 knockdown (Fig 5C). To further explore the role of ENO2 during HCMV infection, we utilized a recently described ENO-specific inhibitor POMHEX, which exhibits a 4-fold higher specificity for ENO2 relative to ENO1 [24]. Treatment with POMHEX substantially inhibited HCMV infection with an ∼85-fold reduction in end-point viral titers relative to control treatment (Fig 5D). We also employed CRISPR to target ENO2 via ENO2-specific CAS9 ribonucleoprotein (RNP) complexes. Electroporation of ENO2-specific CAS9 RNPs successfully reduced HCMV-mediated accumulation of ENO2 (Fig 5E), and had a knockout efficacy of 82% (Fig S3). CRISPR-mediated knockdown of ENO2 was found to significantly reduce the fluorescence associated from a clinically derived HCMV strain, TB40/e, expressing the red fluorescent protein mCherry [22] (Fig 5F). Collectively, these data suggest that induction of ENO2 expression is important for HCMV infection.

**Figure 5.**
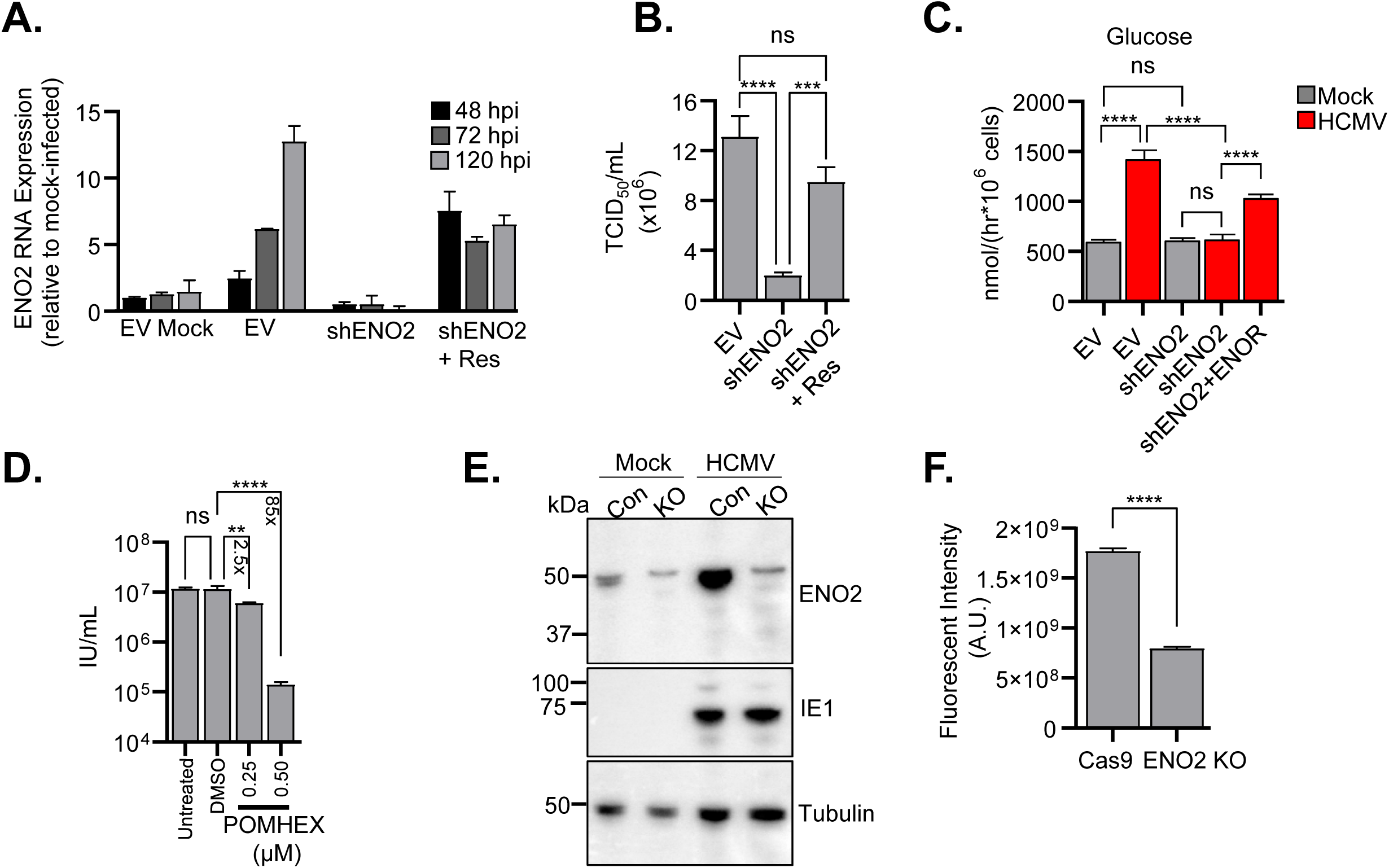
Inhibition of ENO2 attenuates HCMV replication. (A) HFF-hT fibroblasts were transduced with either shRNA targeting Enolase 2 (shENO2), pLKO-empty vector control (EV), or an shRNA-resistant ENO2 (Res). Cells were either Mock or infected with AD169 (MOI 3.0). Total RNA was harvested at the indicated time points. RT-qPCR was performed using ENO2 or GAPDH specific primers. Signals were normalized to GAPDH levels and compared to mock infected cells 48 hpi. (B) TCID_50_/mL analysis was performed from samples harvested at 120 hpi. (C) Total glucose consumed was determined as in Figure 2G. Values are mean ± SEM (N =3). (D) Confluent, serum-starved HFF-hT were infected with AD169 (MOI 3). After inoculation, media were changed and either no drug, DMSO (0.05%), or 0.25 or 0.5 μM POMHEX (0.05% DMSO) were added. Infections were allowed to proceed for 120 hours, at which point virus was taken and IU/mL was determined. Values are mean ± SEM (N ≥12). (E) Polyclonal HFF-hT ENO2 KO cells were successful generated by electroporation of CRISPR RNPs. HFF-hT were electroporated with Cas9 alone without a guide RNA was used as a control. (F) ENO2KO or Cas9 cell lines were plated in 384 well plates and infected at low MOI (0.05) with a TB40-mCherry-GFP virus. mCherry fluorescence was quantified as a readout of viral growth. Values are mean ± SEM (N ≥185). ns, not significant; **, P<0.01; ***, P<0.001; ****, P<0.0001.

### Pharmacological inhibition of ENO2 disrupts HCMV-mediated increases in pyrimidine and pyrimidine-sugar metabolites

To further assess how ENO2 contributes to HCMV-metabolic remodeling, we analyzed metabolite levels upon treatment with the ENO2 inhibitor, POMHEX (Fig 6). Similar to the observations with shENO2 treatment, POMHEX reduced the size of phosphoenolpyruvate pools, and increased fructose-6-phosphate and fructose-bisphosphate pools (Tables S3 & S4, Fig S4). Further, both shENO2 transduction and POMHEX treatment drastically increased the levels of sedoheptulose-7-phosphate, and led to large reductions in pyrimidine and pyrimidine sugar metabolites levels such as N-carbamoyl-L-aspartate, UTP, and UDP-Glucose (Fig 6). While the similarities in the responses to shENO2 transduction and POMHEX treatment were notable, some differences were observed, e.g. shENO2 increased PRPP levels during infection (Fig 3B), while POMHEX treatment reduced PRPP pools (Fig 6). Further, in some cases, POMHEX - induced changes occurred in both mock and HCMV infected cells, e.g., fructose-bisphosphate and UDP-glucose (Fig 6). However, collectively, these data recapitulate the importance of ENO2 in HCMV-mediated modulation of host cell metabolism.

**Figure 6.**
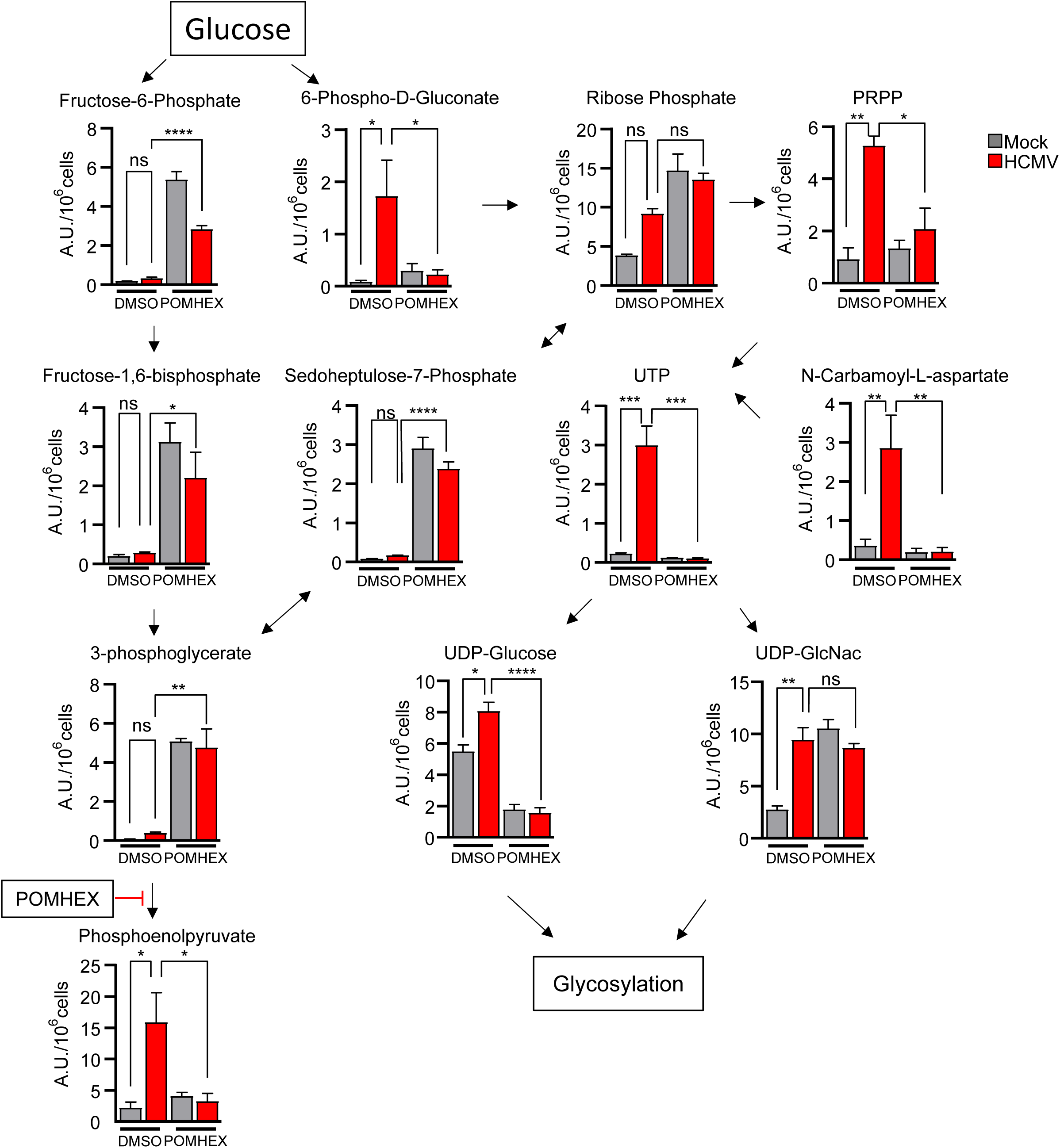
Pharmacological inhibition of ENO2 alters HCMV-mediated metabolic changes. HFF-hT fibroblasts were treated with DMSO (0.05%) or POMHEX (500 nM, 0.05% DMSO) and either Mock or infected with AD169 (MOI 3.0). 48 hpi, metabolic activity was quenched, metabolites extracted, and MS analysis performed. Values are mean ± SEM (N = 3). ns, not significant; *, P<0.05; **, P< 0.01; ****, P<0.0001

To further explore the role of ENO2 in UDP-sugar biosynthesis, we measured the rate of ^13^C-glucose labeling of UDP-sugar pools upon inhibition of ENO2. Inhibition of ENO2 substantially reduced cellular UDP-glucose concentrations, and decreased the ^13^C-labeling rate of UDP-glucose (Fig 7A & 7B), consistent with reduced UDP-glucose biosynthesis. The UDP-glucose pool turned over relatively quickly (Fig S5A), with the MZ+6 isotopologue being the most abundant at all time points (Fig S5B). In contrast to UDP-glucose, the UDP-GlcNac pools in infected cells did not change significantly upon ENO2 inhibition (Fig 7C). UDP-GlcNac labeling was substantially slower than UDP-glucose (Fig 7B & D), but similar to UDP-glucose, the MZ+6 isotopologue was the most prevalent labeled form (Fig S6B). While ENO2 inhibition did not impact infected cell UDP-GlcNac pool sizes, it substantially slowed the labeling of UPD-GlcNac (Fig 7D, Fig S6A), consistent with decreased biosynthesis upon ENO2 inhibition. Taken together, these results indicate that ENO2 is important for UDP-glucose and UDP-GlcNac turnover.

**Figure 7.**
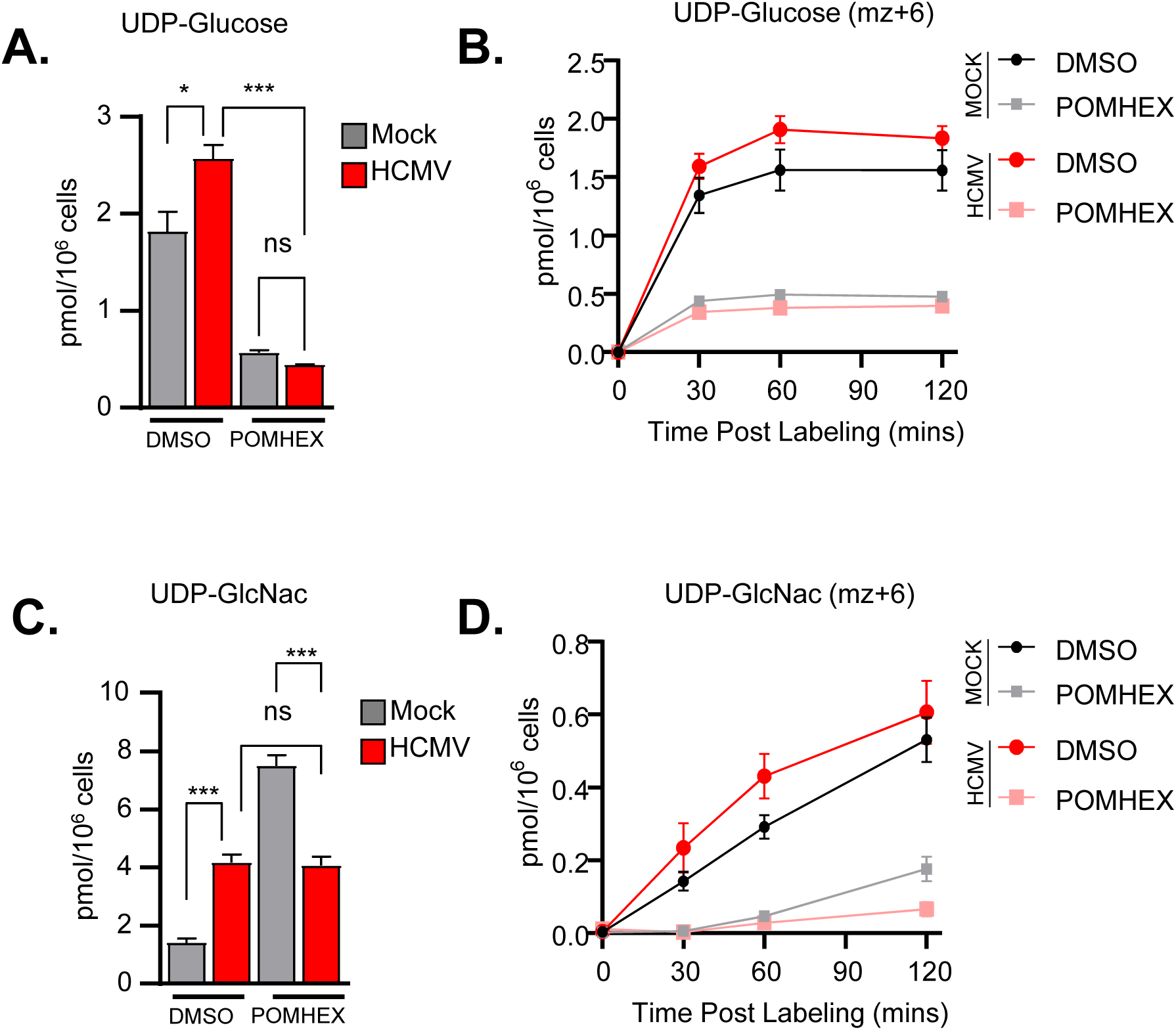
Pharmacological inhibition of ENO2 reduces ^13^C-glucose-mediated labeling of UDP-sugar metabolites. HFF-hT fibroblasts were treated with DMSO (0.05%) or POMHEX (500 nM, 0.05% DMSO) and either Mock or infected with AD169 (MOI 3.0). At 48 hpi, media were switched to U- ^13^C -Glucose and incubated for 0, 30, 60, or 120 minutes before quenching and extraction of metabolites.

### ENO2 is important for the production of infectious HCMV particles and the accumulation of the HCMV envelope glycoprotein B

To investigate how ENO2 might be impacting viral production, we analyzed the accumulation of viral proteins representative of the three stages of viral lifecycle (IE1 = Immediate Early, U_L_26 = Early, pp28 = Late). As shown in Fig 8A, shRNA-mediated reduction in ENO2 expression did not impact the accumulation of viral IE1, U_L_26, or pp28 proteins. Further, loss of ENO2 did not inhibit viral DNA accumulation (Fig 8B). Since the loss of ENO2 did not impact viral DNA accumulation, we reasoned that loss of ENO2 may have a role in particle infectivity. To test this, we quantified packaged viral genomes via a DNase protection assay of partially purified virions. We found that similar levels of packaged, DNase-resistant genomes were produced by control and shENO2 knockdown cells (Fig 8C). However, these virions exhibited an approximately 10-fold reduced infectious titer (Fig 8D), suggesting that many of the genomes produced in shENO2 transduced cells were packaged in defective particles. Literature estimates indicate that HCMV contains one infectious particle for every 100-DNA containing particle [31, 32]. We found that control cells produced 0.863 infectious virions per 100 DNA containing particles, i.e., close to literature estimates (Fig 8E). In contrast, shENO2 cells produced 0.0635 infectious virions per 100 DNA containing particles, an ∼14-fold loss of viral infectivity (Fig 8E). These data indicate that ENO2 is important at a late stage of infection that impacts the production of infectious particles.

**Figure 8.**
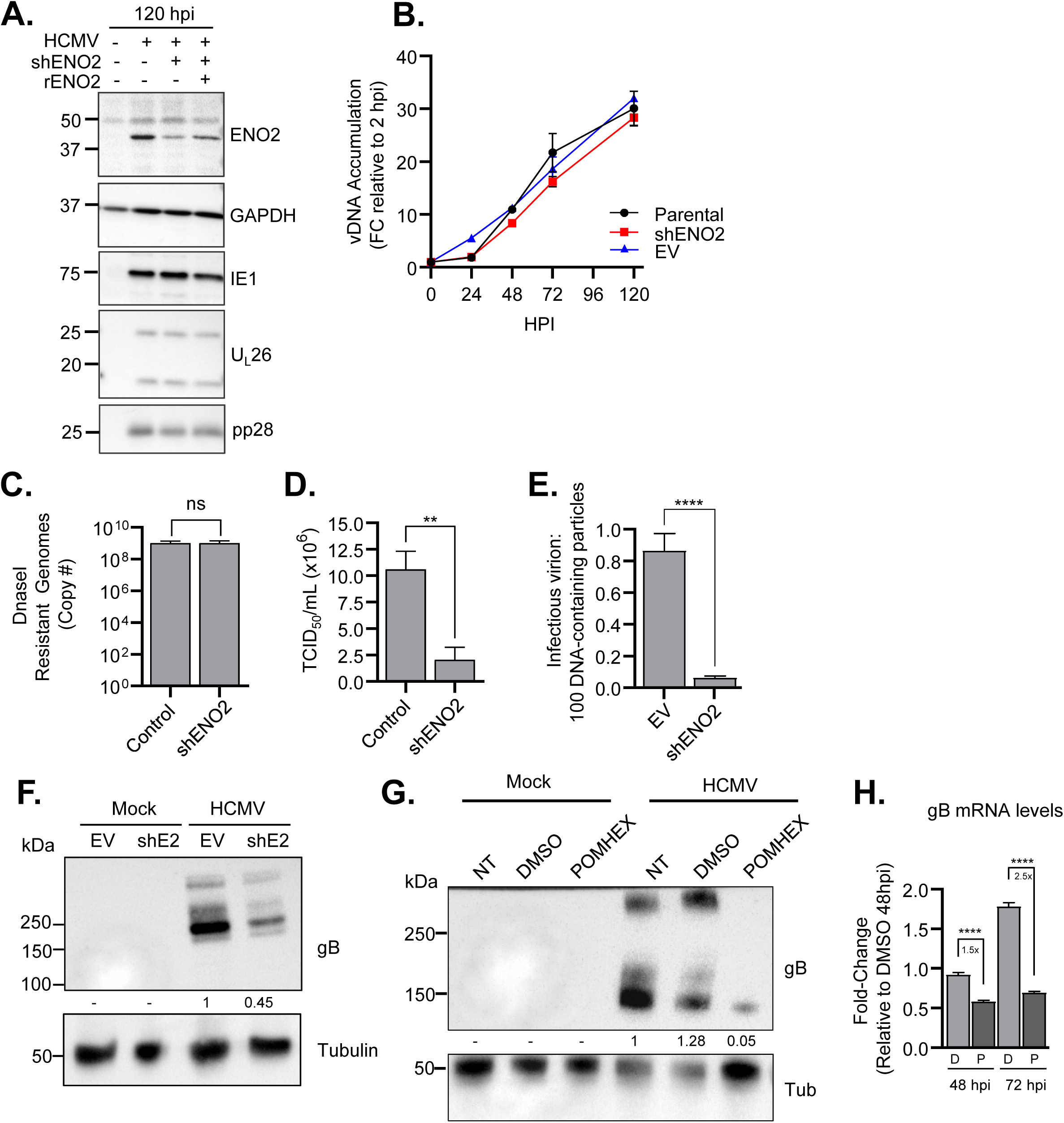
Inhibition of ENO2 attenuates HCMV virion infectivity. (A & B) To determine the impact of ENO2 on the viral lifecycle, HFF-hT were infected with AD169 (MOI 3.0). At indicated time points, total protein and total viral DNA were collected. (A) SDS-PAGE and Western blot analysis of representative viral genes and (B) RT-qPCR analysis of viral DNA accumulation using IE1 specific primers. (C) DNase protected genomes were determined by treating concentrated viral particles with DNase followed by viral DNA extraction. RT-qPCR using IE1 specific primers was used to quantify total viral genomes by comparing to standard curve. (D) TCID_50_/mL was determined from virus collected before DNase treatment. (E) The infectious virion to DNA ratio was determined by dividing the data in C & D, and plotted. (F & G) To determine the impact of ENO2 inhibition on glycoprotein B (gB) during (F) shENO2 or (G) POMHEX treatment, HFF-hTs were AD169 (MOI 3.0) and protein harvested 96 hpi. SDS-PAGE and Western analysis was performed using gB specific antibodies. Protein band intensities are shown relative to EV or NT controls. (H) HFF-hT fibroblasts were either DMSO (D) or POMHEX (P) treated. Cells were infected with AD169 (MOI 3.0). Total RNA was harvested at indicated time points. RT-qPCR was performed using gB or GAPDH specific primers. Signals were normalized to GAPDH levels and compared to 48 hpi DMSO treated samples. Values are mean ± SEM (N = 3). ns, not significant; **, P< 0.01; ****, P<0.0001.

**Figure 9.**
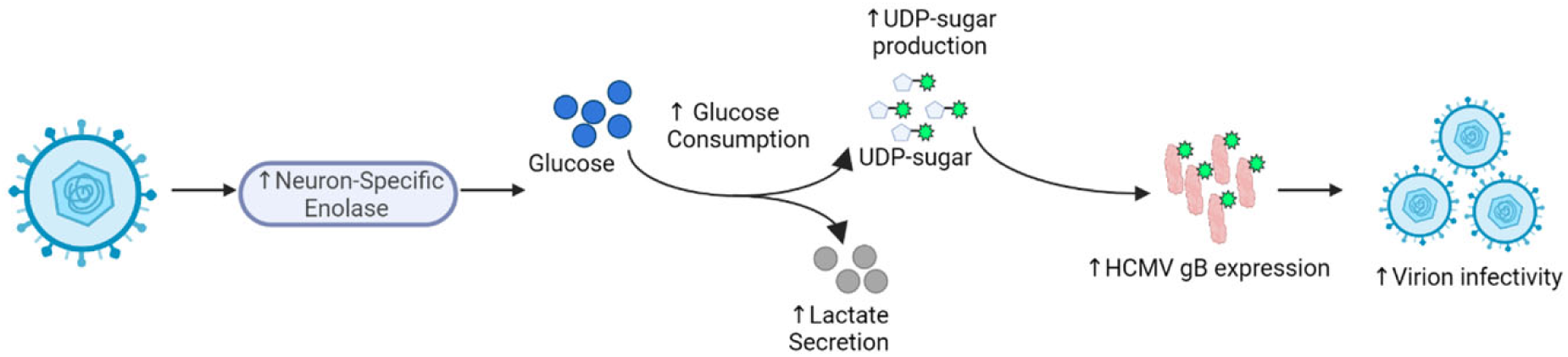
Model of HCMV harnessing neuronal enolase (ENO2) to support infection. HCMV induces the expression of Enolase 2 to increase glycolysis and UDP-sugar pools, which are important for gB expression and virion infectivity.

Previously we had found that pyrimidine biosynthesis is important for the glycosylation of viral but not host proteins [14]. Given the impact of ENO2 on virion infectivity (Fig 8E) and on pyrimidine and UDP-sugar pools (Fig 3A & 6), we hypothesized that ENO2 may be involved in the glycosylation of viral proteins. To investigate this possibility, we analyzed the impact of ENO2 knockdown or inhibition on the accumulation of the viral gB envelope protein (gB). shENO2 treatment resulted in ∼50% reduction in gB accumulation, whereas POMHEX treatment resulted in a 25-fold reduction in gB levels (Fig 8F & 8G). POMHEX also reduced gB mRNA levels (Fig 8H), although the reduction in viral gB mRNA upon POMHEX treatment (∼1-3-fold) was substantially less than what was observed for the impact on gB protein (∼25-fold). These results are consistent with a post-transcriptional impact on gB expression, which would be predicted if a reduction in the availability of glycosylation precursors was responsible for limiting properly folded gB protein accumulation.

## Discussion

Viruses modulate cellular metabolism to support their replication (reviewed in [33]). However, the mechanisms responsible and how these metabolic changes specifically contribute to infection is less clear. We globally assessed how HCMV impacts host gene expression and found that HCMV infection induces a neuronal gene signature (Fig 1). Further, we show that one of the genes induced, the neuronal glycolytic enzyme ENO2, was required for HCMV-mediated glycolytic activation (Fig 2 & 3). Surprisingly, ENO2 was also required for the induction of pyrimidine and pyrimidine sugar pools that are necessary for glycosylation reactions (Fig 3). Inhibition of ENO2, either pharmacologically or with shRNA, attenuated the viral induction of these pools and inhibited the accumulation of the gB glycoprotein (Fig 8F&G). Furthermore, targeting ENO2 resulted in the production of a much higher proportion of defective particles (Fig 8E). Our data suggest a model in which HCMV infection harnesses a tissue atypical neuronal isoform to drive glycolytic and pyrimidine sugar activation to support the production of infectious virions, highlighting a novel facet of HCMV-mediated metabolic modulation.

Humans encode three enolase isozymes, the ubiquitously expressed α-Enolase (ENO1), the muscle specific β-Enolase (ENO3), and neuronal specific γ-Enolase (ENO2 or NSE). Enolase functions as a homo- or heterodimeric complex. The only combination that has not been identified is a βγ dimer, likely due to the differences in tissue expression of each isozyme [34]. The human enolases are ∼80% identical and ∼90% similar and their sequences are highly conserved across species [35]. Many questions remain about how the tissue-specific enolases differ with respect to the contributions made to the tissues in which they are predominantly expressed. The enolases are not thought to be major glycolytic regulators and the step they catalyze, the interconversion of 2-phosphoglycolate to phosphoenolpyruvate, is highly reversible [36, 37]. Despite this, evidence is mounting that the molecular assembly of multienzyme complexes can substantially impact pathway activity [38, 39], highlighting a way in which an enzyme like enolase could be modulating glycolytic activity. Analysis of glycolytic macromolecular enzyme complexes in infected versus uninfected cells could shed new light on these possibilities.

We found that inhibition of ENO2 impacted cellular metabolism in unexpected ways. ENO2 inhibition, whether pharmacological or shRNA-mediated, decreased the product of the rate-determining step of pyrimidine biosynthesis, N-carbomyl-L-aspartate, and attenuated the accumulation UPD-sugar metabolites (Figs 3A & 6). This was surprising given the relative lack of proximity between enolase and the reactions associated with either pyrimidine or UDP-sugar biosynthesis. While the mechanisms through which ENO2 could modulate these distal metabolic pathways remain to be elucidated, there is increasing evidence for metabolites and the enzymes responsible for their production coordinating metabolic responses from afar. Such an example exists for PEP, the product of the enolase reaction. PEP has been found to inhibit Ca^2+^ uptake by the sarcoplasmic/ER Ca2+-ATPase (SERCA) [40, 41]. Recent advances in identifying protein-metabolite interactions could help elucidate novel interactions i.e. how ENO2-mediated disruption of metabolite levels impacts other proteins [42]. With respect to our results, it remains to be determined how ENO2 or PEP could impact the production of N-carbomyl-L-aspartate, but elucidating the mechanisms of distal metabolic coordination will significantly increase our understanding of metabolic regulation in normal physiology and disease.

While enolases have critical metabolic functions, it has also been observed that they have roles outside of their traditional glycolytic activity. ENO1 localizes primarily to the cytoplasm, but can also translocate to the cell membrane where it acts as a plasminogen receptor [34]. Further, a truncated ENO1 isoform called myc-binding protein 1 (MBP1) can localize to the nucleus, where it acts as a tumor-suppressor that can restrict the expression of the c-Myc oncogene [43]. In contrast to ENO1, the localization of ENO2 and ENO3 have not be extensively studied; however, ENO2 and ENO3 have also both been described to have potentially non-glycolytic roles. For example, ENO3 is important for muscle cell development and regeneration, which appears to be independent of its glycolytic activity [44]. In addition, ENO2 has been shown to have neurotrophic and neuroprotective effects on neurons. A c-terminal peptide of ENO2 is sufficient to facilitate these effects in neurons [45]. Some studies suggest an ENO1/ENO2 dimer is expressed on the surface of neuronal cells [46]. Additionally, ENO2 associates with GAPDH and other proteins to form a trans-plasma-membrane oxioreductase complex that acts as an extracellular sensor to signal external oxidative stress [47]. Given these reports, we currently cannot rule out the possibility that ENO2’s contribution to pyrimidine and pyrimidine-sugar biosynthesis and the normal production of infectious particles is independent of its glycolytic activity.

Our results indicate that inhibition of ENO2 does not substantially impact the early times of infection, but rather, decreases the accumulation the gB glycoprotein and increases the production of defective viral particles. We have previously found that HCMV infection upregulates pyrimidine and nucleotide sugar biosynthesis for the production of UPD-sugars necessary for the glycosylation of viral proteins, and which if inhibited resulted in a similar phenotype, i.e., mostly normal viral protein and DNA production, but a defect in envelope protein accumulation and decreased infectivity [14]. With respect to our current ENO2 results, the importance of UDP-sugar levels for the normal accumulation of glycoprotein B (gB) agree with these findings. However, substantial questions remain about how HCMV infection modulates pyrimidine and pyrimidine-sugar biosynthesis. For example, to what extent do viral factors specifically modulate pyrimidine and pyrimidine-sugar activities to support viral protein glycosylation?

We targeted ENO2 in two ways, with shRNA and with a novel Enolase inhibitor POMHEX [24]. While there was substantial overlap in the impact of these two treatments on both metabolite pools and viral infection, there were significant differences, e.g., on some glycolytic pools (compare Figs 3 & 6), or on the magnitude of the changes induced to gB expression, or production of infectious virions (Figs 5 & 8). These differences could reflect a more complete inhibition of ENO2 upon POMHEX treatment relative to shRNA-mediated targeting, although potential off-target effects of either the shRNA or POMEX cannot be ruled out.

Our RNA-seq data indicate that HCMV infection induces a neuronal gene expression signature, which includes the expression of ENO2. To our knowledge, this has not been described during HCMV infection. Our data indicate that part of this transcriptional rewiring is responsible for supporting HCMV-mediated metabolic remodeling. At this time, it is unclear how the induction of this neuronal-like environment might contribute to other facets of infection. We anticipate that there will be other neuronal genes induced that are found to be important to infection, yet it remains to be seen. Notably, a recent report suggests that Kaposi’s Sarcoma-Associated Herpesvirus (KSHV) induces a neuroendocrine transcriptional environment during latent infection of endothelial cells [48]. While the situations are quite different, e.g. lytic versus latent infection, taken together, these results suggest the possibilities that certain viruses have evolved to harness tissue-specific epigenetic programs to support their infection.

Increasingly, viruses are found to modulate cellular metabolism to support their infection (reviewed in [33, 49]). Here, our data suggest that a virus can harness tissue atypical gene programs to metabolically remodel the host cell. These findings open up wide ranging possibilities with respect to the mechanisms through which viruses can modify the cellular metabolic environment to procure the metabolic resources they require for successful infection. Future experiments are necessary to elucidate whether harnessing atypical gene expression programs is a common viral tactic of viral metabolic reprogramming.

## ACKNOWLEDGMENTS

The work was supported by NIH grants AI127370 and AI50698 to J.M. I.M. was supported by T32 training grants (AI118689 and GM68411), and by a pre-doctoral fellowship (AI157283-01). IRS was supported by a pre-doctoral fellowship from the American Heart Association.

---

**Supplemental Figure 1.**
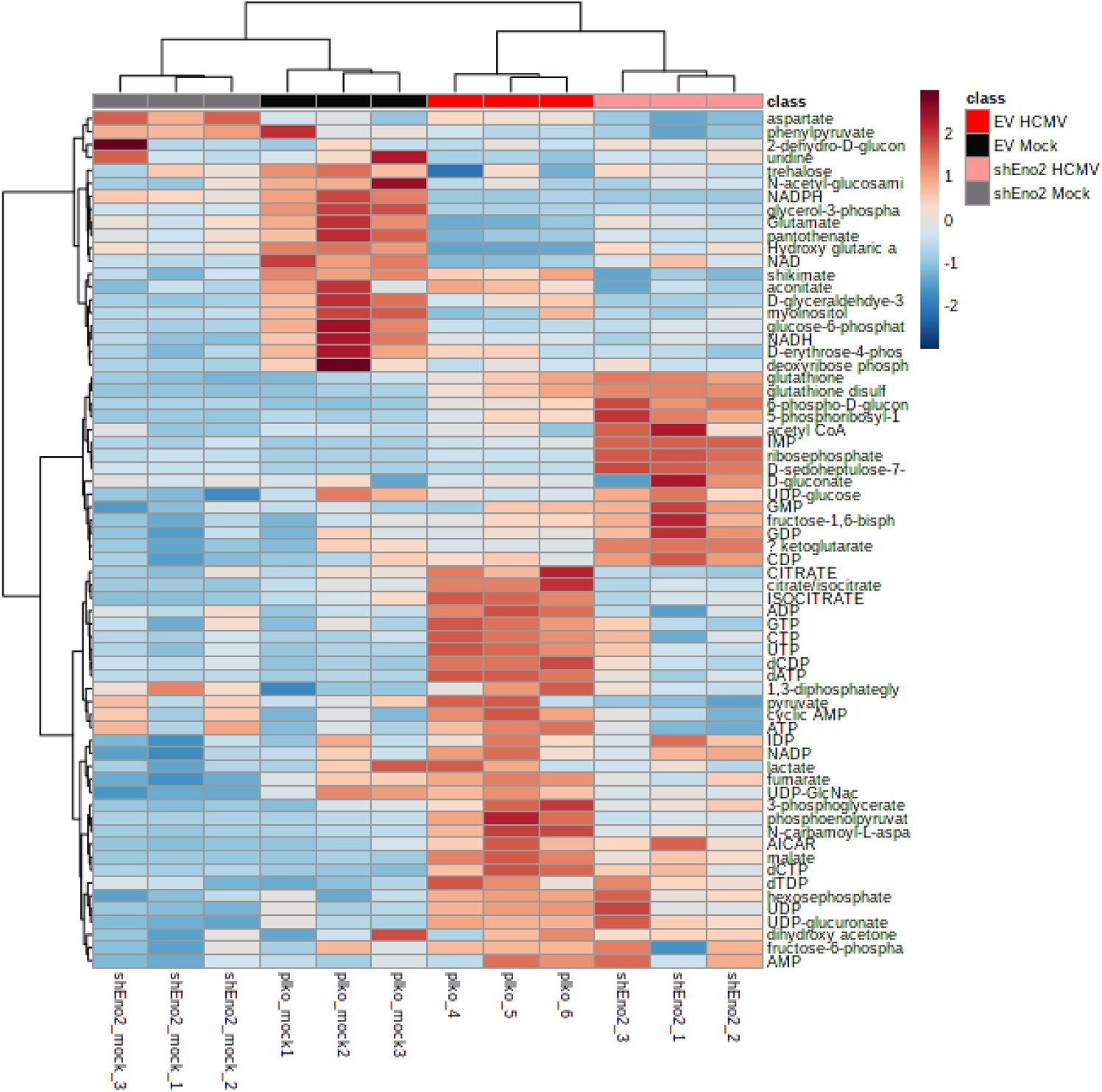
Analysis of shEno2 metabolic changes. Related to Figure 2. Heat map of metabolites identified by MS after shENO2 treatment. Hierarchal clustering showing relationship between Mock (Black/Gray) and HCMV infected (Red/Pink) samples.

**Supplemental Figure 2.**
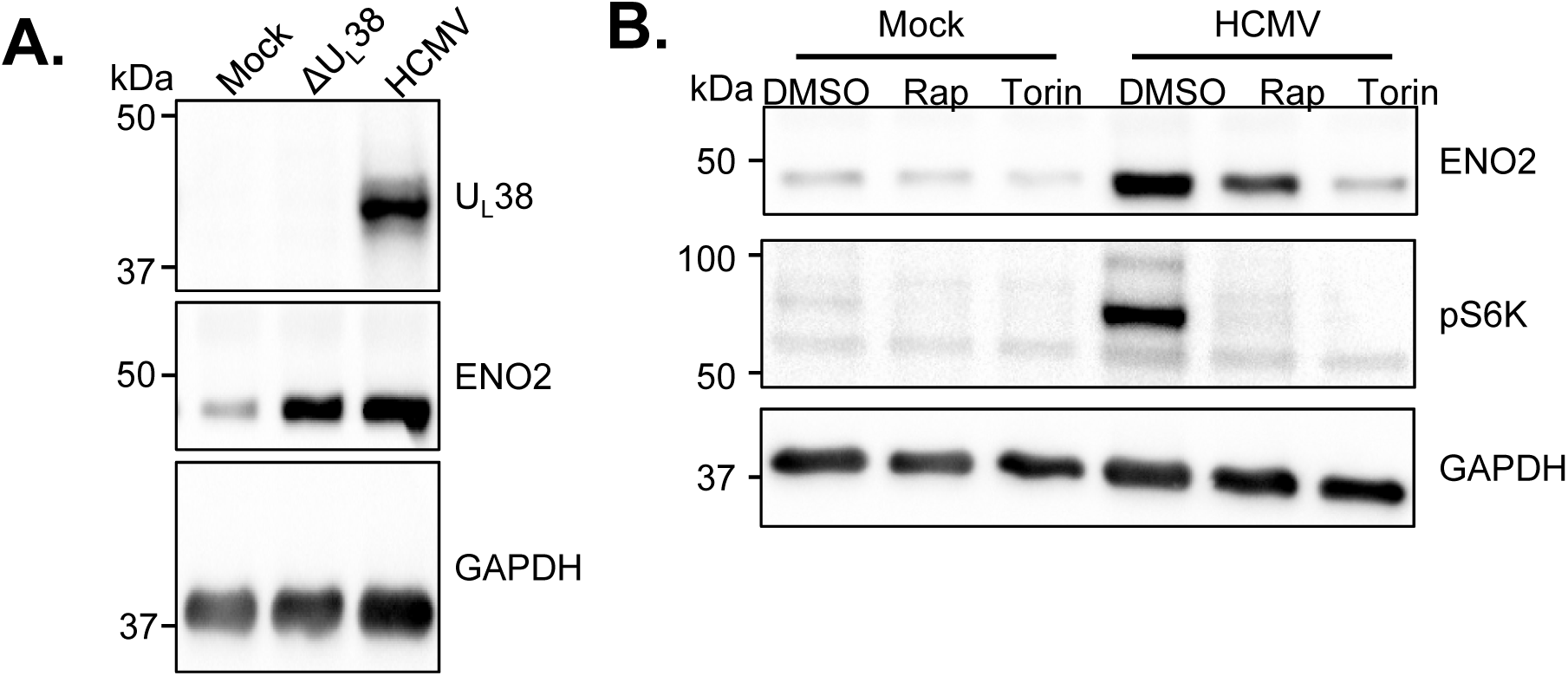
HCMV protein U_L_38 and mTOR activation are required for Eno2 expression in MRC5 Fibroblasts. Related to figure 4. (A) MRC-5 fibroblasts were either mock, AD169 infected (MOI 3.0), or AD169ΔUL38 infected (MOI 3.0). Total protein was collected 48 hpi and western blot analysis performed with the indicated antibodies. (B) MRC-5 fibroblasts were either mock or AD169 infected (MOI 3.0) and treated with Rapamycin, Torin, or DMSO. Total protein was collected 48 hpi and western blot analysis performed with the indicated antibodies.

**Supplemental Figure 3.**
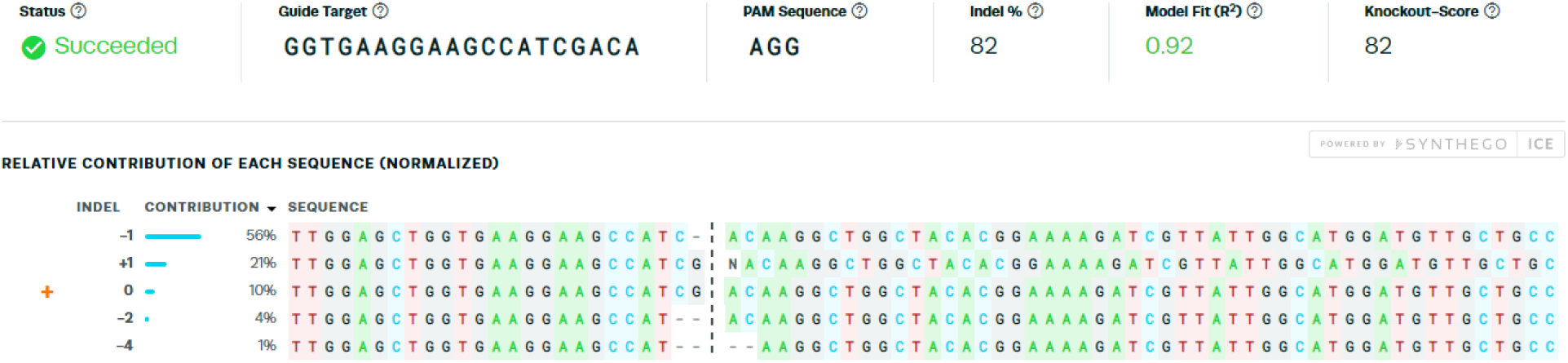
Synthego ICE Analysis of ENO2 CRISPR Knockout polyclonal cell lines. Related to figure 5. HFF-hTs edited by CRIPSR-Cas9 to knockout Enolase 2. RNPs containing Cas9 and duplexed gRNA:tracrRNA were electroporated into HFF-hTs. Once cell lines recovered, total DNA was extracted and ENO2 genomic DNA was amplified. This amplicon was then sequenced. The sequencing reads were then deconvoluted by Synthego’s Inference of CRISPR Edits (ICE) analysis.

**Supplemental Figure 4.**
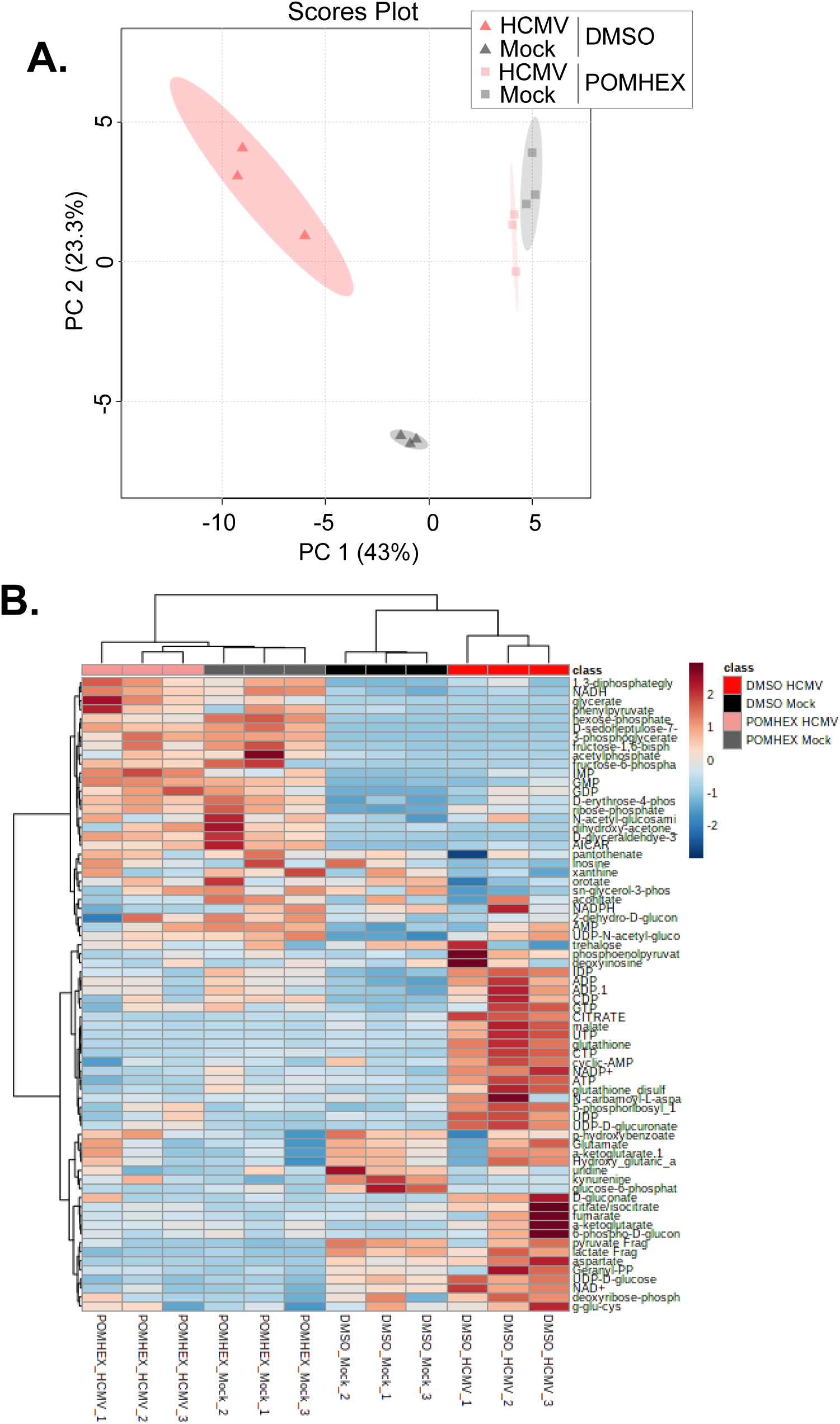
Analysis of POMHEX metabolic changes. Related to Figure 6. (A) PCA analysis and (B) Heat map of metabolites identified by MS Hierarchal clustering showing relationship between Mock (Black/Gray) and HCMV infected (Red/Pink) samples.

**Supplemental Figure 5.**
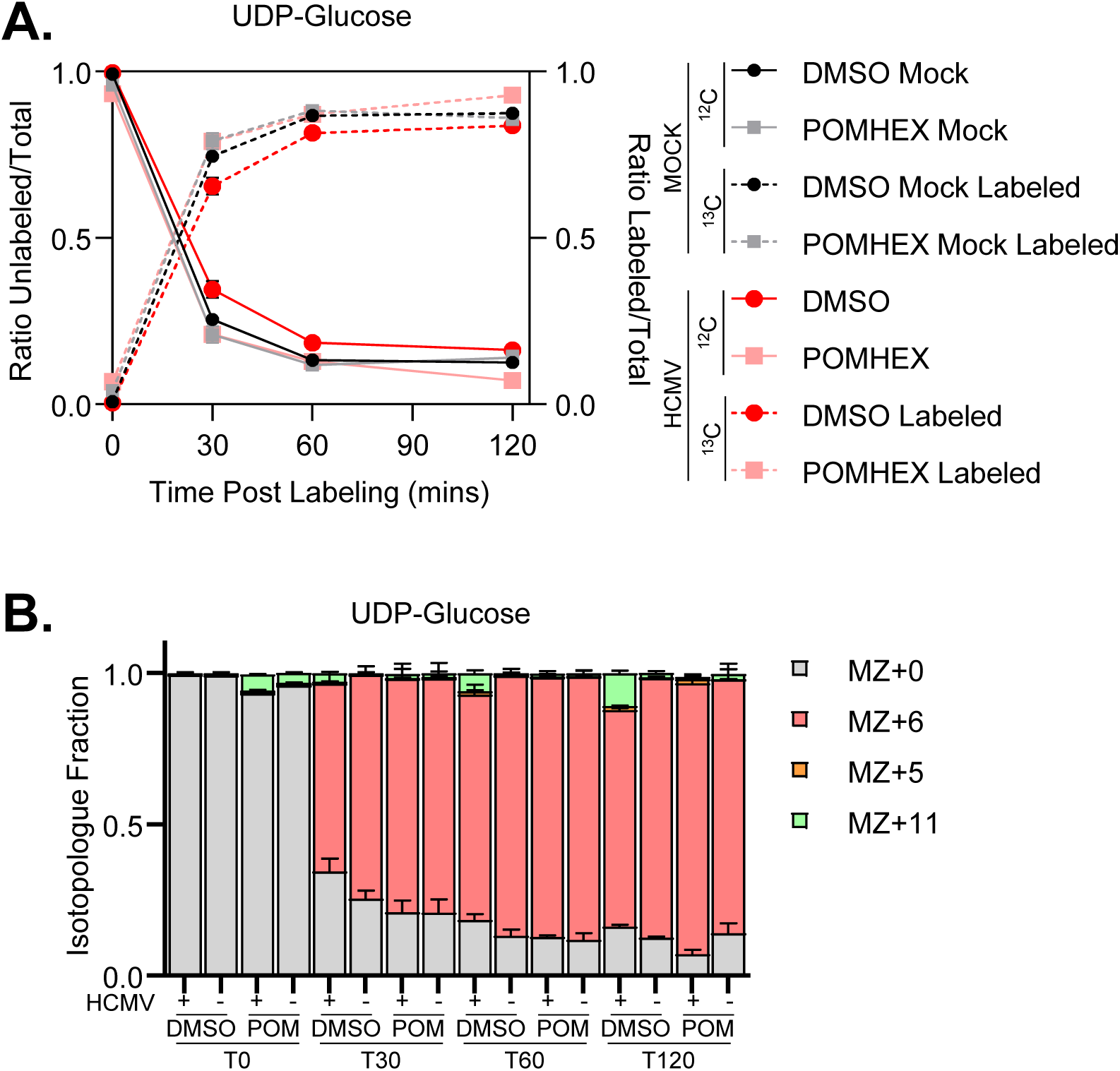
Rapid turn-over of UDP-Glucose metabolic pool in HFF-hT. Related to Figure 7 A&B. (A) Ratio of unlabeled and labeled UDP-Glucose metabolites are shown as a function of time post addition of the ^13^C label. (B) Breakdown of isotopologues by sample. ^12^C (MZ+0) shown in gray, signal derived from the direct addition of ^13^C-Gluose to UDP (MZ+6) shown in pink, ^13^C-glucose conversion to ribose of UDP (MZ+5) shown in orange, and fully labeled UDP-Glucose where the ribose ring and glucose moieties are fully labeled shown (MZ+11) in green.

**Supplemental Figure 6.**
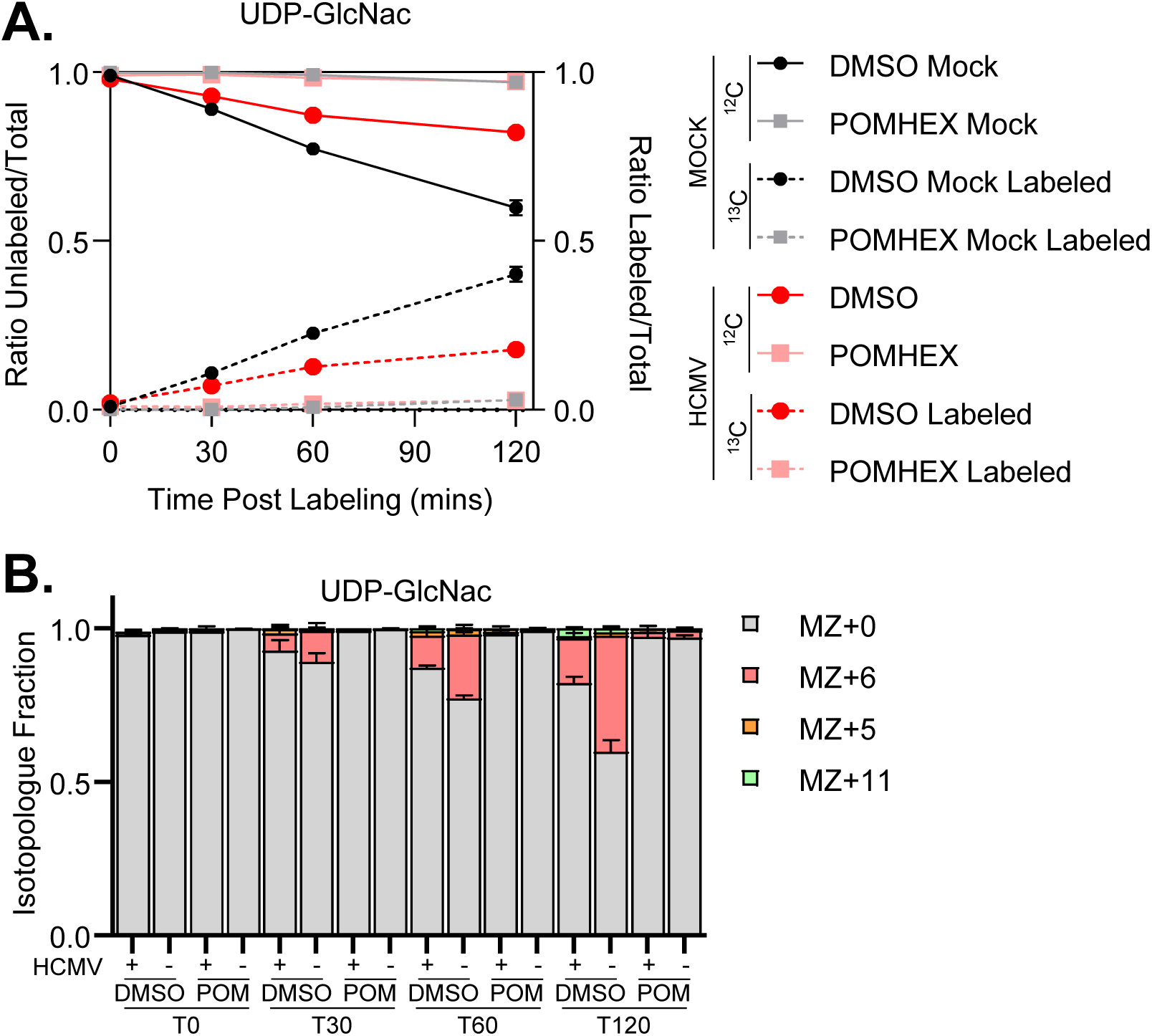
Slow turn-over of UDP-GlcNac metabolic pool in HFF-hT. Related to Figure 7 C&D. (A) Ratio of unlabeled and labeled UDP-GlcNac metabolites are shown as a function of time post addition of the ^13^C label. (B) Breakdown of isotopologues by sample. ^12^C (MZ+0) shown in gray, signal derived from the conversion of ^13^C-Gluose to n-acetylglucosamine and added to UDP (MZ+6) shown in pink, ^13^C-glucose conversion to ribose of UDP (MZ+5) shown in orange, and fully labeled UDP-GlcNac where the ribose ring and glucose moieties are fully labeled shown (MZ+11) in green.

